# Goal-Directed Learning is Multidimensional and Accompanied by Diverse and Widespread Changes in Neocortical Signaling

**DOI:** 10.1101/2023.02.13.528412

**Authors:** Krista Marrero, Krithiga Aruljothi, Christian Delgadillo, Sarah Kabbara, Lovleen Swatch, Edward Zagha

## Abstract

New tasks are often learned in stages with each stage reflecting a different learning challenge. Accordingly, each learning stage is likely mediated by distinct neuronal processes. And yet, most rodent studies of the neuronal correlates of goal-directed learning focus on individual outcome measures and individual brain regions. Here, we longitudinally studied mice from naïve to expert performance in a head-fixed, operant conditioning whisker discrimination task. In addition to tracking the primary behavioral outcome of stimulus discrimination, we tracked and compared an array of object-based and temporal-based behavioral measures. These behavioral analyses identify multiple, partially overlapping learning stages in this task, consistent with initial response implementation, early stimulus-response generalization, and late response inhibition. To begin to understand the neuronal foundations of these learning processes, we performed widefield Ca^2+^ imaging of dorsal neocortex throughout learning and correlated behavioral measures with neuronal activity. We found distinct and widespread correlations between neocortical activation patterns and various behavioral measures. For example, improvements in sensory discrimination correlated with target stimulus evoked activations of licking-related cortices along with distractor stimulus evoked global cortical suppression. Our study reveals multidimensional learning for a simple goal-directed learning task and generates hypotheses for the neuronal modulations underlying these various learning processes.

## Introduction

The recent surge in combining rodent behavioral training with longitudinal neuronal recordings provides unprecedented opportunities for revealing the neuronal mechanisms underlying task learning (e.g., (Chen et al., 2015; Hedrick et al., 2024; Huber et al., 2012; Komiyama et al., 2010; Laubach et al., 2000; Makino et al., 2017; Musall et al., 2019; Peters et al., 2014; Poort et al., 2015; Rabinovich et al., 2022)). However, when performing such studies, the investigators must decide which behavioral measures to track and which brain regions to monitor. For example, motor skill learning has traditionally been correlated with neuronal changes in motor cortex (Laubach et al., 2000; Peters et al., 2017), while sensory detection and sensory discrimination learning has traditionally been correlated with neuronal changes in sensory cortex (Chen et al., 2015; Poort et al., 2015). And yet, lesioning and recording studies suggest that multiple brain regions contribute to learning, even for simple goal-directed tasks (Koralek et al., 2013; Lashley, 1950; Le Merre et al., 2018; O’Doherty, 2004; Strick et al., 2009; Taylor & Ivry, 2012).

We consider two, non-exclusive explanations for the involvement of multiple brain regions in simple task learning. First, learning across a single objective outcome (e.g., skilled reaching or sensory detection) may be broadly distributed across multiple neuronal networks. Second, operant learning may involve multiple learning stages, with each learning stage having its own objective outcome and corresponding neuronal mechanisms. In support of this second explanation, a recent modeling study of rodent operant conditioning demonstrated different learning rates for different task outcome measures (for example, response bias vs perceptual sensitivity) (Ashwood et al., 2020). Additionally, perturbation and recording studies have demonstrated that the contributions of specific brain regions are limited to specific learning phases (Hwang et al., 2021; Kawai et al., 2015). These studies suggest diverse learning stages in rodent operant tasks. However, for the multitude of goal-directed learning tasks currently employed, the structure of their learning stages and the outcome objectives of each stage remains unknown.

Here, we leverage a classic skill learning framework from cognitive psychology. Fitts and Posner (1967) proposed three learning stages to skill acquisition. In the cognitive stage the performer develops an understanding of the sensorimotor action (the ‘executive program’) required for task success; in the associative stage the performer practices and refines the initial sensorimotor action sequence learned in the cognitive stage; in the autonomous stage the skill is acquired and conducted with less cognitive effort than the preceding stages (Fitts & Posner, 1967). It is currently unknown whether this Fitts and Posner framework of learning stages is useful for rodent operant tasks.

In this study, we tracked multiple behavioral measures across learning of 52 mice in a Go/NoGo operant whisker discrimination task (respond to target stimuli, ignore distractor stimuli) (Aruljothi et al., 2020). We tracked both object-based (post-stimulus response rates and measures of detection/discrimination) and temporal-based (pre-stimulus sampling and post-stimulus reaction time) behavioral outcome measures. Using unbiased methods, we determined whether changes in these behavioral measures are distributed in a manner suggestive of learning stages. Furthermore, in a subset of mice, we performed longitudinal widefield Ca^2+^ imaging of dorsal neocortex from naïve to expert task performance. From these combined behavior-neuronal datasets we constructed maps of the neuronal activation patterns that correlate with changes in various behavioral measures. We sought to determine whether changes in different behavioral measures correlate with changes in the same neuronal regions (low dimensional) or different neuronal regions (high dimensional). From these analyses we propose two general conclusions: 1) even for a simple operant task, mouse learning occurs in stages that resembles the Fitts and Posner framework, and 2) learning across these stages involves diverse (high dimensional) and widespread changes in neocortical signaling. These conclusions provide the foundation for deeper understandings of the neuronal mechanisms underlying goal-directed task learning.

## Materials and Methods

### Animal subjects

Experimental protocols have been approved by the IACUC of University of California, Riverside. The behavioral dataset used here includes studies that have been previously reported (Aruljothi et al., 2020; Marrero et al., 2022; Zareian et al., 2023; Zareian et al., 2021; Zhang & Zagha, 2023). Wild type (C57BL/6J, JAX #000664; BALB/cByJ, JAX #001026), transgenic (Snap25-2A-GCaMP6s-D, JAX #025111; Thy1-ChR2-YFP, JAX #007612; VGAT-ChR2-EYFP, JAX #014548), and virus-injected adult male and female mice were included in the behavioral learning data. Mice were not under transgenic manipulation (e.g., optogenetic activation) during behavioral training. Mice were maintained in a 12-h light/dark cycle and behavioral training occurred during the light cycle. Mice started behavioral training after a minimum of a 4-day recovery period from surgery.

### Animal behavior

Training stages, metrics of learning, and criterion for expert performance in the Go/NoGo whisker discrimination task were as previously reported (Aruljothi et al., 2020; Marrero et al., 2022; Zareian et al., 2021). Briefly, head-fixed and water deprived mice learned on customized behavioral rigs. Behavioral data was collected using Arduino and custom MATLAB scripts (MathWorks, MA). Target and distractor paddles were positioned symmetrically within the mouse whisker fields bilaterally. Mice reported stimulus detection by licking a centrally located lick port, positioned under the snout. Mice progressed through classical conditioning (2-5 days) and operant conditioning (2-5 days) stages before entering the full task. Data presented in this paper are exclusively from the full task, in which the task structure did not change through training.

In the full task we implemented a variable intertrial interval (ITI) before a stimulus presentation to discourage anticipatory responding. Licking within the ITI was punished by resetting the ITI, randomly selected from a negative exponential distribution between 5.5 and 9.5 s. Target stimuli and distractor stimuli were presented on a probabilistic schedule, 20% and 80% respectively. Following the correct rejection of a distractor stimulus, a shortened ITI was randomly selected from a negative exponential between 0.2 and 1.9 s followed by a target trial (such that the longest wait time for a target trial was 14.6 s). We also implemented a short lockout period (200 ms or 300 ms) after a stimulus presentation. Responding during the lockout ended the current trial and initiated a full ITI. Lick bouts were considered spontaneous (sampling) if they occurred within the ITI. Post-stimulus licking responses were considered hits if they occurred after a target whisker stimulus and during the response window (after the lockout). Post-stimulus licking responses were considered false alarms (FA) or premature (Preme) if they occurred after a distractor whisker stimulus or during the lockout window, respectively. Hits (responses to target stimuli) were rewarded with ∼5 μL of water, correct rejections (not responding to distractor stimuli) and correct withholdings (not responding during the catch trial) were rewarded with a shortened intertrial-interval (ITI) and a subsequent target trial. Only trials within periods of task engagement were analyzed, with ‘task engagement’ defined as a minimum block of 10 minutes without a pause in licking greater than 60 seconds. If more than one engagement period was identified within a session, the longest engagement period was chosen for the subsequent analyses. Spontaneous rates per session were calculated as the percentage of trials in which the mouse responded within 1 second before an impending stimulus. Response rates per session were calculated as the percentage of response trial outcome types (hit, false alarm, premature) across all stimulus trials. Measures based on signal detection theory were calculated,

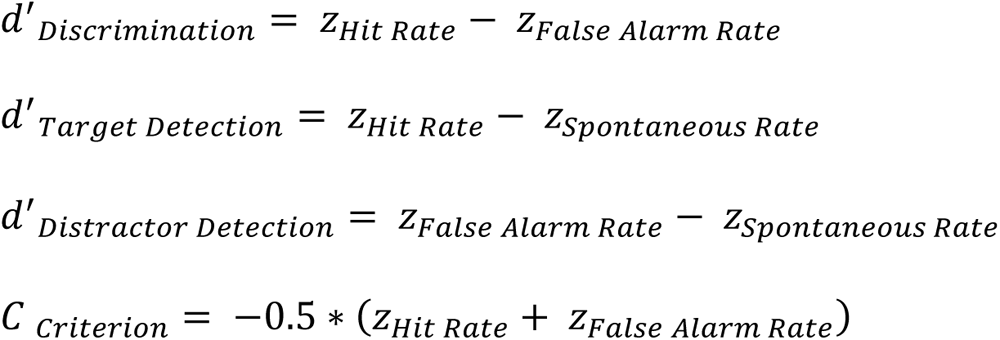

where z is the inverse normal distribution function.

### Structure of the behavioral training data

For interlick interval (ILI) analyses, ‘sampling AUC’ was determined as the percent area under the histogram curve of spontaneous lick bouts between zero and the minimum ITI per session. For most sessions, this was between zero and 5.5 seconds. ‘Waiting AUC’ was determined as the percent area under the histogram curve of post-stimulus responses between the minimum ITI per session and the maximum possible wait time for a target stimulus (14.6 s). The post-stimulus lockout period was 0.2 seconds for 49 mice and 0.3 seconds for 3 mice. Hit and false alarm reaction times were bounded by the lockout period and the subsequent 1 second response window. Data are reported as mean ± standard error of the mean (SEM) unless otherwise noted. Cumulative probabilities per session, per tercile, per tercile per mouse, and by performance level were determined by cumulative distribution functions (CDFs), the normalized cumulative sum of ILI or RT distributions. Because interlick and reaction time distributions were not normally distributed, the median values (instead of means) are reported.

### Data inclusion criteria

For behavioral measures, we examined 120 mice with potential learning data but included only the 52 mice who met all criteria described below. First, mice were included if they learned the task, as defined by discrimination performance (d’ between hit rate and false alarm rate) greater than one for three consecutive days. Second, mice were included if they learned in less than three weeks of training. Third, the data required a minimum of seven training sessions with a maximum 7-day gap between sessions. No more than seven sessions of consecutive expert behavior were analyzed.

For tercile and quintile learning data per mouse, each session was binned according to the exclusive integer value of total sessions divided by three and five, respectively. For learning data binned by performance, each session was defined as naïve (d’<0.8), intermediate (0.8<d’<1.2), and expert (d’>1.2). This allowed for comparison of time in training (tercile or quintile) versus learning stage (performance).

### Behavioral learning analyses

Analyses were performed using standard MATLAB scripts. Linear regression analyses were used to determine slopes of behavioral outcome measures across sessions per mouse. To assess behavioral measure dynamics (monotonic, biphasic, non-phasic) we performed one-way ANOVA on quintile-binned data from individual behavioral measures per mouse. ‘Monotonic’ was based on significant differences across quintiles, with the first (decreasing) or last (increasing) quintile value being the largest. ‘Biphasic’ was based on significant differences across quintiles, with the middle (2, 3, or 4) quintile value being the largest. ‘Non-phasic’ was based on non-significant differences across quintiles. Unsupervised clustering by k-means analyses was performed using the *kmeans* function in MATLAB on the quintile data averaged across mice and amplitude normalized. Number of clusters was determined by knee point detection. Rate analyses (Δvelocity/Δquintile) for determining transition acceleration or deceleration were performed on normalized and rectified (monotonically increasing) quintile data.

### Widefield Ca^2+^ imaging

Widefield Ca^2+^ imaging was performed as previously described (Aruljothi et al., 2020). However, for these studies mice were imaged daily from naïve through expert task performance in the full task. Imaging was acquired from GCaMP6s express-ing Snap25-2A-GCaMP6s-D mice (JAX #025111), which express GCaMP6s in both excitatory and inhibitory neurons throughout the brain (Madisen et al., 2015). Head-post implantation and skull preparation were performed as described in (Aruljothi et al., 2020). The widefield imaging system consists of a Macroscope IIa (RedShirt Imaging, beam diverter removed), 75 mm inverted lens with 0.7x magnification and 16 mm working distance. Optical excitation was provided by a mounted 470 nm LED (Thorlabs M470L3) dispersed through a collimating lens (Thorlabs ACL2520-A), band-pass filtered (Chroma ET480/40x), and directed through the macroscope using a dichroic mirror (Chroma T510lpxrxt). Emission light was band-pass filtered (Chroma ET535/50m) and acquired by an RT sCMOS camera (SPOT Imaging). This configuration yields a field of view of 7 mm x 5.8 mm, 41 μm pixel diameter, 12-bit depth, 10 Hz image capture. TIF image sequences were imported to MATLAB for further processing.

### Sliding Window Normalization and Trial-Based Neuronal Activity

Trial-based imaging was initiated 1 s before stimulus onset and continued through the 0.2 s lockout window and the subsequent 1 s response window, totaling 2.2 s per trial. F_n_(i,j,f) represents the fluorescence (F) of each pixel at row i and column j in frame f for trial n. To standardize fluorescence values, we initially computed the local mean using a sliding window approach, calculated every 2 seconds with a window size of +/-200 seconds [F_SW_(i,j,n)]. Subsequently, normalized fluorescence values were determined for each pixel in each frame following the methodology outlined by (Salkoff et al., 2020):

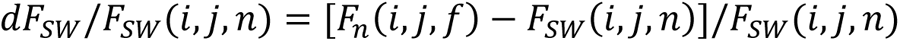

To generate average movies, we first sorted trials based on stimulus type (target and distractor). Subsequently, we computed the mean pixel-wise activity across frames within each trial category. Imaging data across sessions were aligned by bregma. Resulting activity maps were rigidly referenced to the Allen Brain Institute’s common coordinates framework (CCF) (Wang et al., 2020) and other functionally defined regions of interest (Chen et al., 2017; Ferezou et al., 2007; Mayrhofer et al., 2019).

### Quantification of post-stimulus activation

Post-stimulus activation was quantified by computing the neurometric d’ (Britten et al., 1992) comparing prestimulus fluorescence (stimulus absent) to post-stimulus fluorescence (stimulus present), as previously performed (Aruljothi et al., 2020; Marrero et al., 2022). Neurometric d’ was calculated separately for target and distractor trials. Prestimulus and post-stimulus fluorescence histograms were converted into receiver operating characteristic (ROC) curves, and the area under the curve (AUC) was then translated into d’, serving as the neurometric measure:

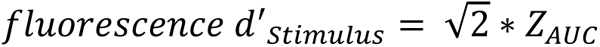

These neurometric d’ measures per pixel across sessions were used for behavioral-neuronal correlation analyses (using the *corrcoef* function in MATLAB). For example, hit rate per session was correlated with the neurometric d’ on target trials per session, individually for all pixels. For data shown in this study, the frame 0.2-0.3 s post-stimulus was used as the measure of stimulus-evoked activation, which includes both post-stimulus and pre-response signaling.

### Statistics

All analyses, including statistics, were performed using standard and custom MATLAB scripts and visualized using CorelDRAW. For one-way analysis of variance (ANOVA) statistical tests, Tukey-Kramer post-hoc multiple comparisons were generated using function *multcompare* in MATLAB. All repeated measures ANOVA statistical tests were conducted in Prism with Tukey-Kramer post-hoc multiple comparisons between measures, within subject, and between sex as applicable. Significance was assigned according to p-values <0.05. For visualization, asterisks are shown according to orders of p-values: *p<0.05, **p<0.005, ***p<0.0005. For imaging analyses, statistical tests were used to determine post-stimulus activation differences between expert vs naïve sessions (unpaired t-test) or behavioral-neuronal correlations significantly different from zero across mice (one-sample t-test). Instead of a significance threshold, we display ‘significance maps’ as the -log_10_ of the calculated p-value per pixel.

## Results

### Operant whisker discrimination task structure and primary outcome measures

In this Go/NoGo task, water-restricted mice learn to selectively respond to target paddle deflections in one whisker field (hit, rewarded with water delivery) and ignore distractor paddle deflections in the opposite whisker field (correct rejection) (Figure 1A-B). During the variable intertrial interval (ITI), mice were required to withhold responding, as spontaneous responding caused a reset of the ITI. Following the ITI, target and distractor stimuli were presented probabilistically (see methods). Overall, mice were required to wait (withhold licking) across a duration of minimum ITI of 5.5 s to maximum 14.6 s to be presented with a target stimulus and opportunity for reward. Additionally, we implemented a minimum 200 ms lockout period after a stimulus presentation, through which the mice were required to withhold before responding to obtain a reward.

In this study we present longitudinal behavioral training data from 52 mice. Expert performance in this task is determined solely by target vs distractor stimulus discrimination (discrimination d’ > 1 for three consecutive days). To compare across mice with different training durations, we segmented training sessions into terciles (or quintiles, below).

Discrimination significantly increased across terciles (Figure 1C, one-way ANOVA F(2,573)=167.49, p<0.0005). Across the same training sessions, we observed a decrease in the criterion, indicating an increased tendency to respond across target and distractor trials (Figure 1D, one-way ANOVA F(2,573)=59.47, p<0.0005).

**Figure 1.**
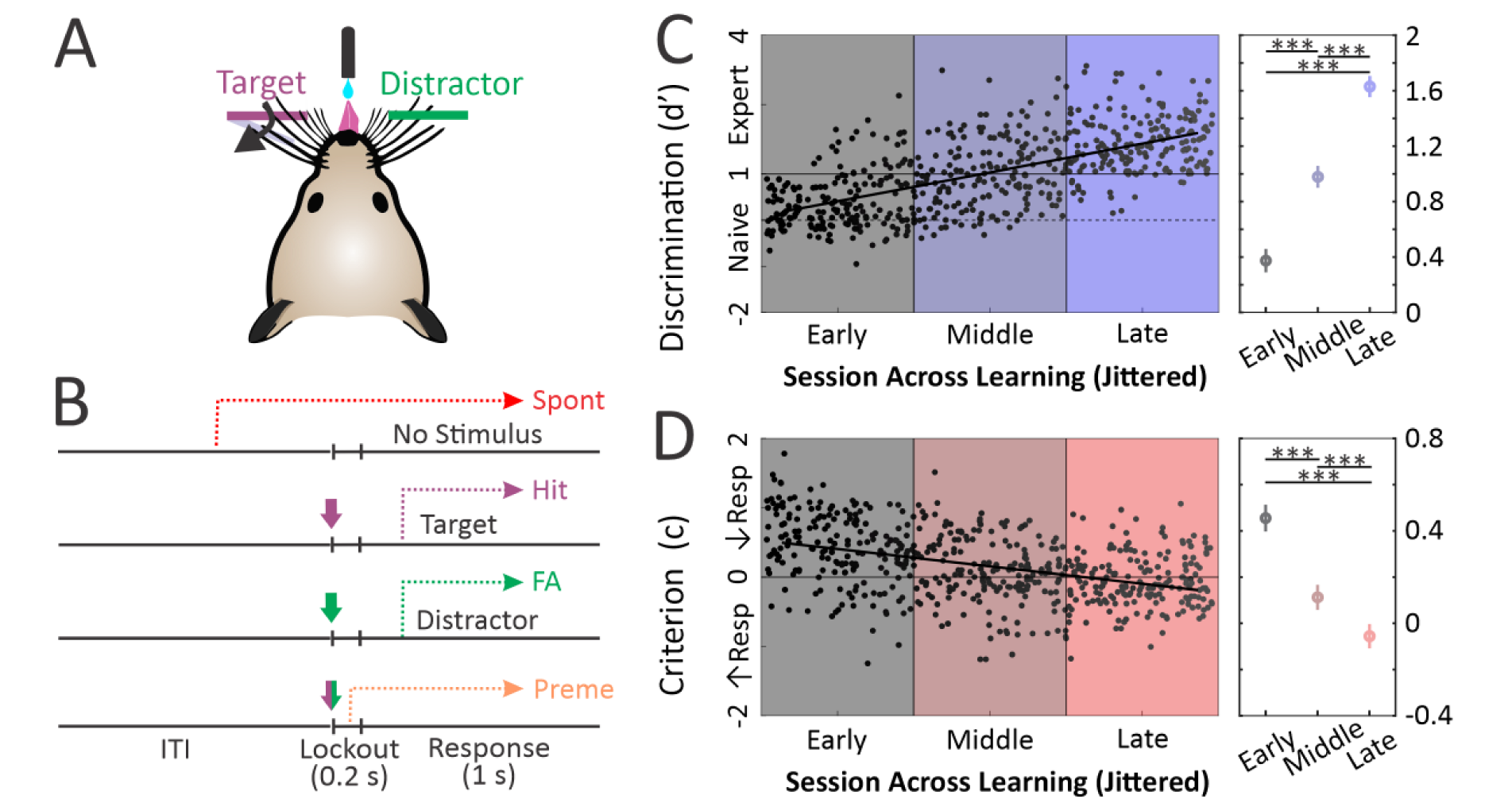
Trial structure and performance. (A) Mice learn to selectively respond to target whisker deflections within one whisker field (fuchsia) and ignore distractor whisker deflections in the opposite whisker field (green). (B) Trial structure and response assignments. A target or distractor stimulus is presented on separate trials. Spontaneous (Spont, red) response during the ITI and before stimulus presentation; Hit (fuchsia) response after a target stimulus during the response window; False alarm (FA, green) response after a distractor stimulus during the response window; Premature (Preme, orange) response after a stimulus during the lockout period. (C) Target vs distractor discrimination performance plotted across all sessions and all mice, segregated according to duration in training (early, middle, late) (D) Same dataset and layout as in [C], but for criterion bias. Shading denotes progression in training by tercile, dark to bright. One-way ANOVA with post-hoc multiple comparisons, *** p<0.0005.

### Quantification of pre-stimulus waiting behavior

Given the strict restrictions on withholding during the ITI, we sought to quantify each mouse’s waiting behavior (Figure 2). First, we distinguished ‘sampling behavior’ and ‘waiting behavior’. Sampling behavior was defined as responding before the minimum ITI (5.5 s), which invariably caused a resetting of the ITI. Waiting behavior was defined as responding after successfully waiting for a stimulus. We investigated sampling and waiting behavior across training (Figure 2A-D). Notably, waiting increased significantly from early to middle terciles and remained elevated, with a nonsignificant increase from middle to late terciles (Figure 2C, one-way ANOVA F(2,573)=22.20, p<0.0005). We normalized sampling and waiting distributions to all responses, generating ‘sampling AUC’ (AUC, area under the curve) and ‘wait AUC’ metrics. Across training, sampling AUC decreased (one-way ANOVA F(2,573)=24.19, p<0.0005) while wait AUC increased (one-way ANOVA F(2,573)=42.15, p<0.0005) (Figure 2D).

**Figure 2.**
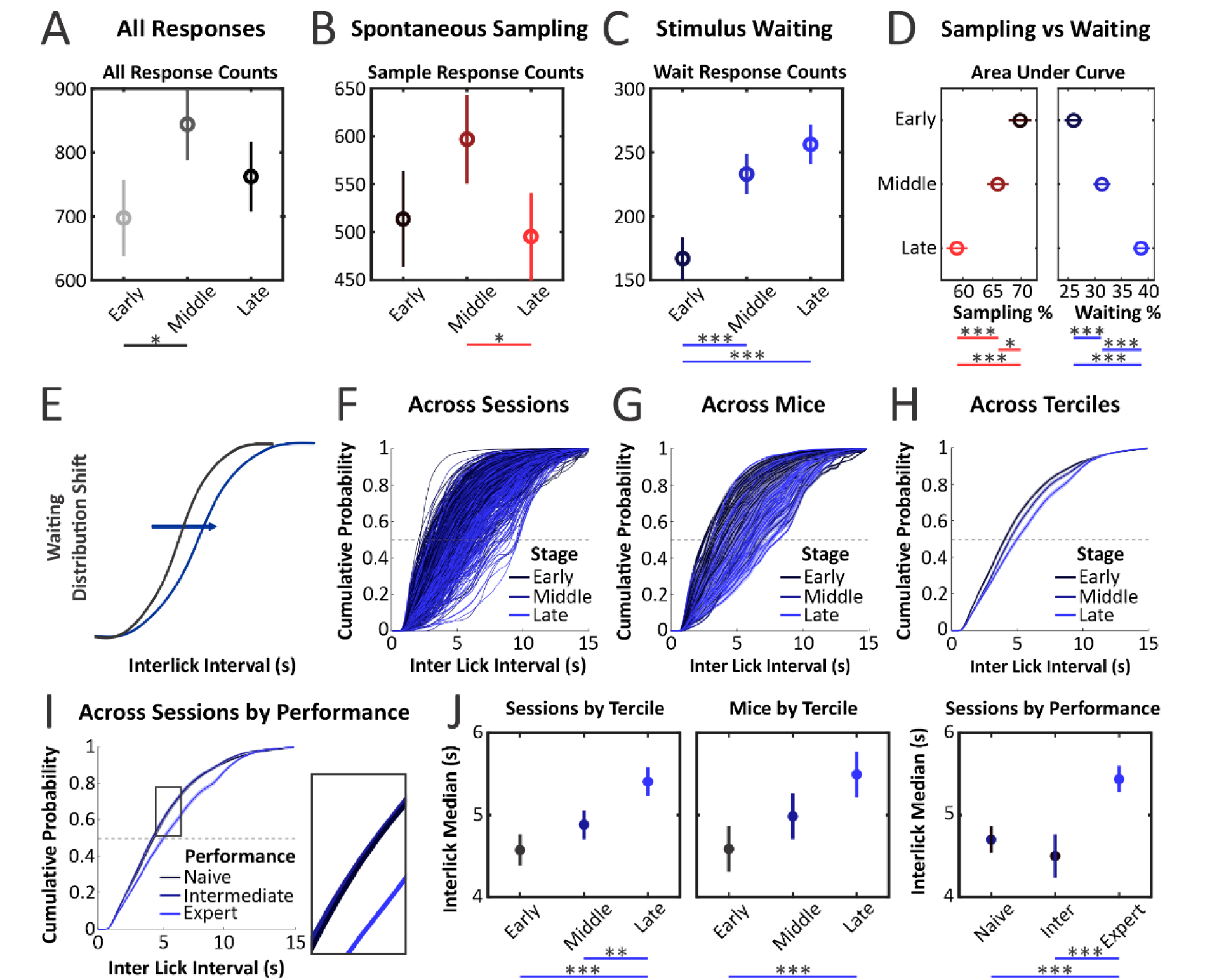
Mice transition from a sampling to waiting strategy before stimulus presentation across learning. (A) Mice increased all responding from early to middle terciles. (B) Mice decreased spontaneous sampling (prestimulus responding) from middle to late terciles. (C) Mice increased stimulus waiting (post-stimulus responding) from early to middle and late terciles. (D) Percent area under the curve (%AUC) per session for spontaneous sampling (red) and stimulus waiting (blue). As a percentage of all responding, spontaneous sampling decreased and stimulus waiting increased across training. (E) Expected cumulative distribution function shift if waiting behavior improves across learning. (F) Cumulative probability distribution function (CDF) curves across all sessions. (G) CDF curves across terciles per mouse. (H) Mean CDF curves across terciles averaged across mice. (I) Mean CDF curves for sessions binned according to discrimination performance; inset shows overlapping of naïve and intermediate performance CDFs. (J) Quantification of median interlick intervals across sessions by tercile training, across mice by tercile training, and across sessions by discrimination performance. [A] Shading denotes progression in training, grey to black; [B-J] shading denotes progression in training, dark to bright; [F-I] dashed line denotes 50% CDF for median value estimation. One-way ANOVA with post-hoc multiple comparisons * p<0.05, ** p<0.005, *** p<0.0005.

We also investigated waiting behavior by the cumulative distribution functions (CDF) of the interlick interval (ILI, time between bouts of licking/responding) distributions (Figure 2E-J). With more effective waiting we would expect a rightward shift of the ILI CDF (Figure 2E).

Consistently, we observed a rightward shift in the ILI distributions with training (Figure 2F-H). In addition to indexing to time in training (terciles), we also plotted ILI CDF curves according to target vs distractor discrimination, for naïve (d’<0.8), intermediate (0.8<d’<1.2), and expert (d’>1.2) task performance (Figure 2I-J). Interestingly, we found that ILI CDF curves during naïve and intermediate performance were similar, yet with a rightward shift during expert sessions (one-way ANOVA F(2,573)=20.34, p<0.0005). These analyses indicate that enhanced ability to wait correlates with expert, but not intermediate or naïve, task performance. Furthermore, they indicate potentially non-uniform learning trajectories for different performance measures, which we explore further below.

### Quantification of post-stimulus timing behavior

Next, we investigated reaction times (RT) post-stimulus (Figure 3). Given the 200 ms lockout period, we were particularly interested in RT convergence around the lockout duration. Responses to all stimuli (Figure 3A), responses after target stimuli during the lockout window (premature, Figure 3B), and responses after target stimuli during the response window (hits, Figure 3C) increased most robustly about the 200 ms lockout duration. To quantify this, we measured the mean RTs across sessions and terciles (Figure 3D). Hit RTs significantly decreased across terciles (one-way ANOVA F(2,569)=100.51, p<0.0005). Notably, target premature RTs significantly *increased* from early to late terciles (one-way ANOVA F(2,545)=5.09, p=0.0065). This convergence of RTs is consistent with a post-stimulus timing strategy to match the reaction time to the lockout duration.

**Figure 3.**
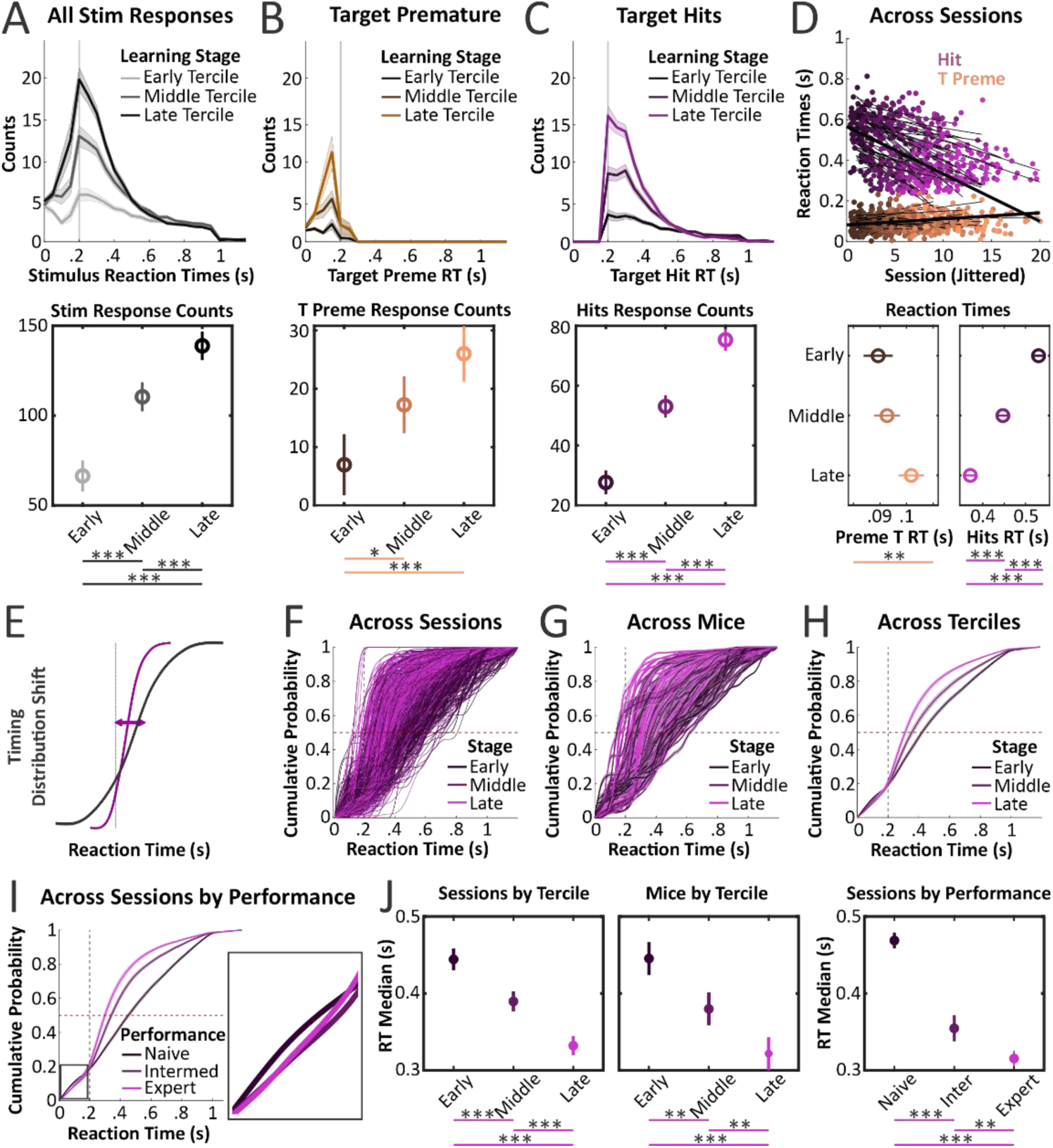
Mice transition from a sampling to timing strategy after stimulus presentation across learning. (A) Mice increased post-stimulus responses across terciles. Vertical line [A-C, E-I] denotes the end of the post-stimulus, pre-response lockout window. (B) Mice increased post-target premature (orange) responses from early to middle and late terciles. (C) Mice increased target hit (purple) responses across terciles. (D) Target stimulus RTs per session for hits and target premature trials with linear fits across sessions per mouse (thin black lines) and mean of linear fits across mice (bold black lines). Target premature RTs increased from early to late terciles whereas hit RTs decreased across all terciles (E) Expected cumulative distribution function shift if timing behavior improves across learning. (F) Cumulative probability distribution function (CDF) curves across all sessions. (G) CDF curves across terciles per mouse. (H) Mean CDF curves across terciles averaged across mice. (I) Mean CDF curves for sessions binned according to discrimination performance; inset shows overlapping of intermediate and expert performance CDFs. (J) Quantification of median RTs across sessions by tercile training, across mice by tercile training, and across sessions by discrimination performance. [A] Shading denotes progression in training, grey to black; [B-J] shading denotes progression in training, dark to bright; [F-I] horizontal dashed line denotes 50% CDF for median value estimation. One-way ANOVA with post-hoc multiple comparisons * p<0.05, ** p<0.005, *** p<0.0005.

As timing behavior improves, we would expect an RT distribution to shift towards the lockout duration (Figure 3E). We investigated the RT CDF and found it to shift left (towards the lockout duration) across sessions, across mice, and across terciles (Figure 3F-H). Similar to waiting behavior, we also plotted RT CDF curves according to target vs distractor discrimination. Interestingly, we found that RT CDF curves during intermediate and expert performance were significantly shifted compared to naïve performance (Figure 3I-J). These analyses indicate that mice do show improvements in timing. Yet unlike waiting behavior, improvements in timing behavior appeared to lead expert sensory discrimination, providing further evidence for non-uniform learning trajectories.

Interestingly, changes in timing behavioral did not generalize across stimulus types. Reponses counts to distractor stimuli false alarms (FAs) did increase from early to middle terciles of training (Figure 4B). However, there were no significant changes in RTs for distractor trials (premature or FA, Figure 4C). Thus, improvements in timing behavior were specific for target stimuli.

**Figure 4.**
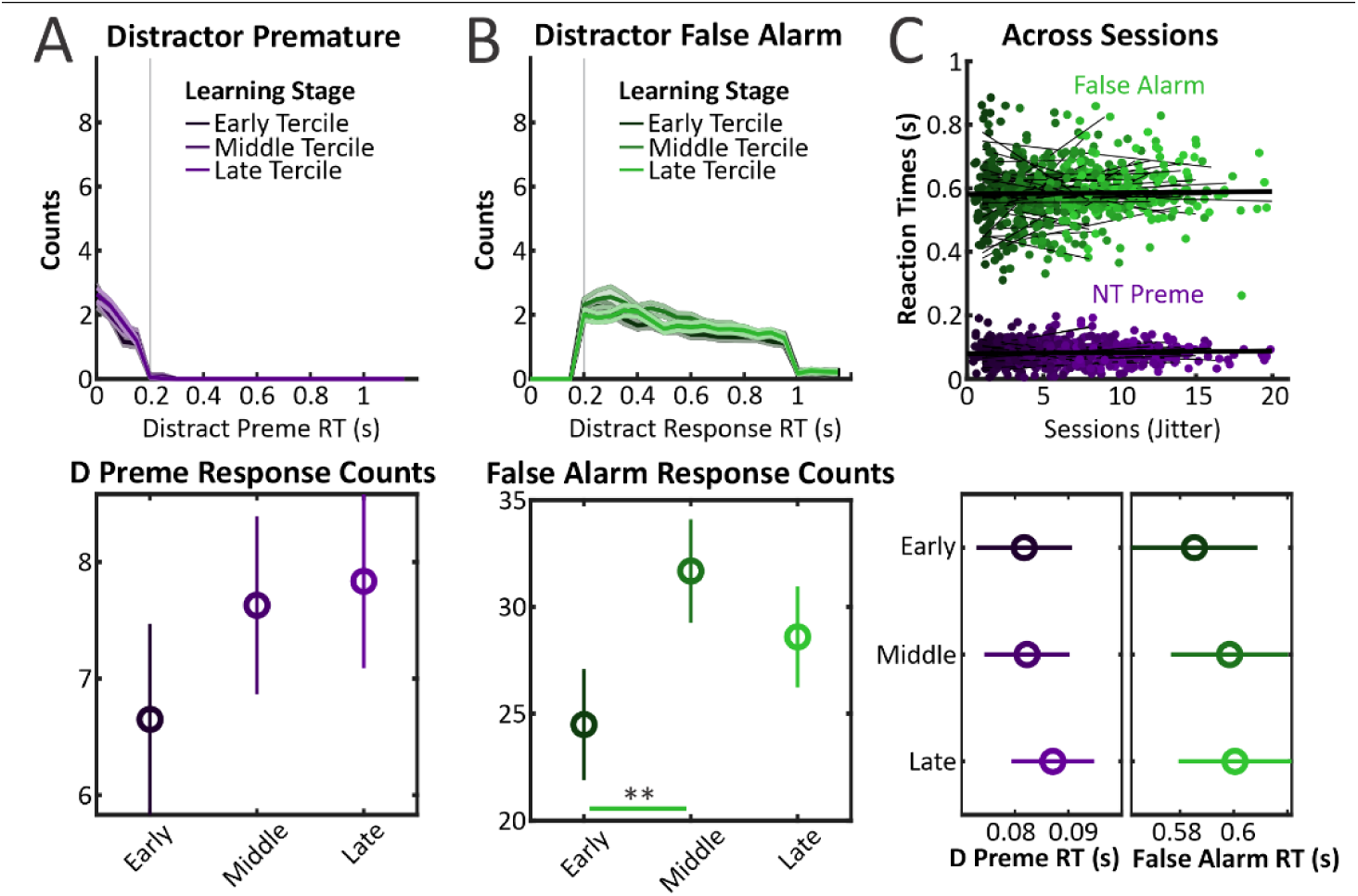
Mice do not demonstrate changes in timing behavior for distractor trials. (A) Distractor premature (purple) response counts did not change across terciles. (B) Distractor false alarm (green) response counts increased from early to middle terciles. (C) Distractor RTs per session for false alarm and premature trials with linear fits across sessions per mouse (thin black lines) and mean of linear fits across mice (bold black lines). Shading denotes progression in training, dark to bright. Note, unlike target RTs, distractor RTs do not change across training. One-way ANOVA with post-hoc multiple comparisons ** p<0.005.

### Comparing behavioral measures averaged across all mice

In Figure 5 we present changes across training for 11 different behavioral measures related to spontaneous and post-stimulus responding. [Note, not all behavioral measures are independent, such as target detection which is the separation between hit rate and spontaneous rate.] Some behavioral measures exhibited monotonic changes with training (increasing: response rates, hit rates, target detection, target vs distractor discrimination, wait AUC, Figure 5A) (decreasing: target RT, criterion, Figure 5B).

**Figure 5.**
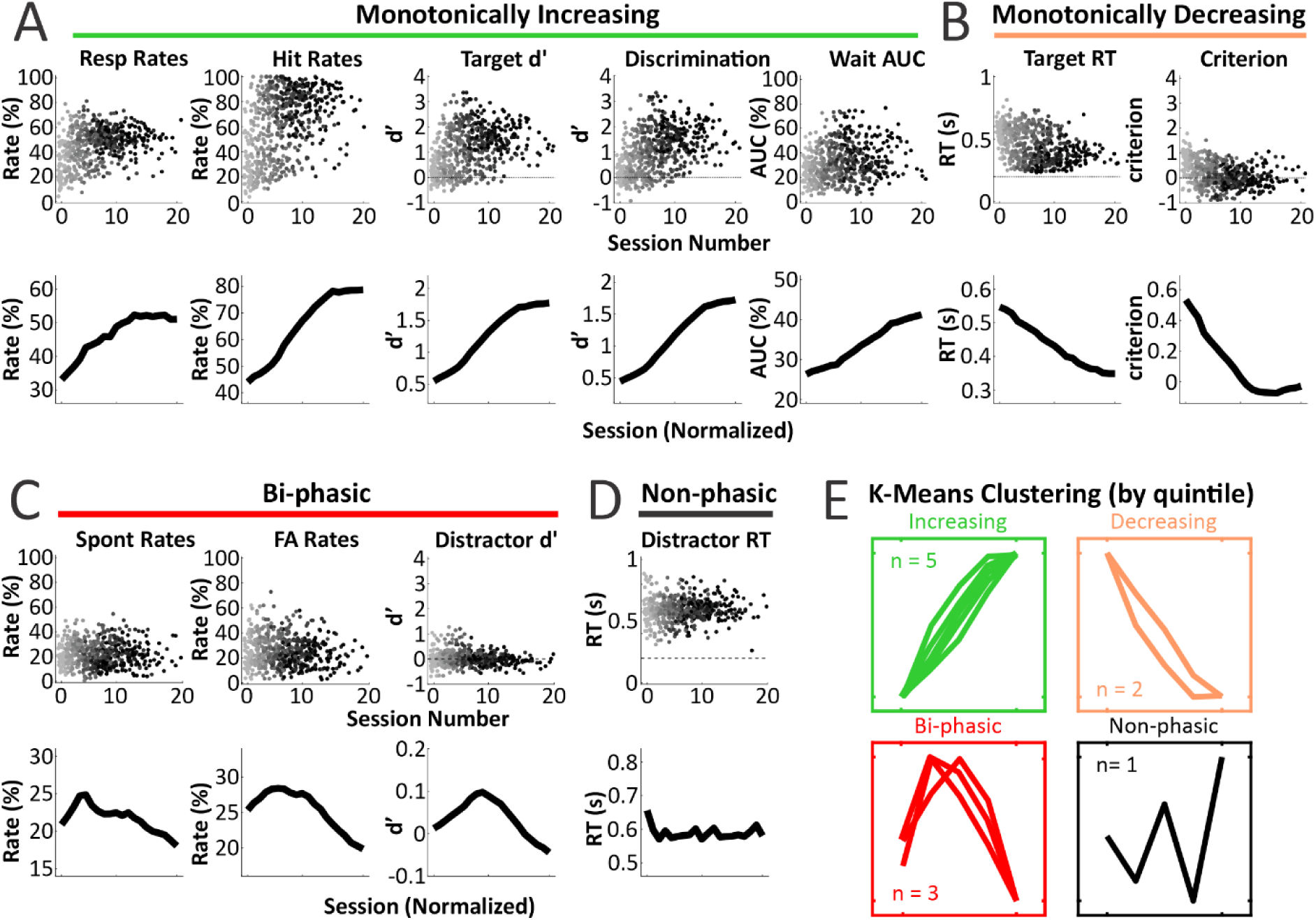
Mice display robust changes in multiple behavioral measures. (A) Monotonically increasing behavioral measures across sessions across mice: response (Resp) rate, hit rate, target detection d’, target vs distractor discrimination d’, wait AUC. (B) Monotonically decreasing behavioral measures: target RT, criterion. (C) Biphasic behavioral measures: spontaneous (Spont) rate, false alarm (FA) rate, distractor detection d’. (D) Non-phasic behavioral measures: distractor RT. Shading denotes progression in training, grey to black. Top rows in [A-D], session numbers are jittered for visualization. Bottom rows in [A-D] display the mean data across mice with normalized training durations. (E) K-means clusters of normalized, quintile-binned behavioral measures averaged across mice, with the same color scheme as in [A-D].

Some behavioral measures did not significantly vary with training (distractor RT, Figure 5D). Perhaps most revealing, other behavioral measures displayed biphasic changes with training (spontaneous rates, false alarm rates, distractor detection, Figure 5C). Notably, all measures with monotonic trajectories contain measures of correct (rewarded) behavior whereas all measures with biphasic trajectories are measures of incorrect (punished) behavior. The biphasic learning trajectories, in particular, suggest multi-staged learning, with generalized responding early in training and withholding of incorrect responding later in training. Unsupervised clustering (k-means, see methods) of the averaged and normalized quintile learning trajectories support these four groupings (monotonically increasing, monotonically decreasing, biphasic, non-phasic) (Figure 5E).

**Figure 6.**
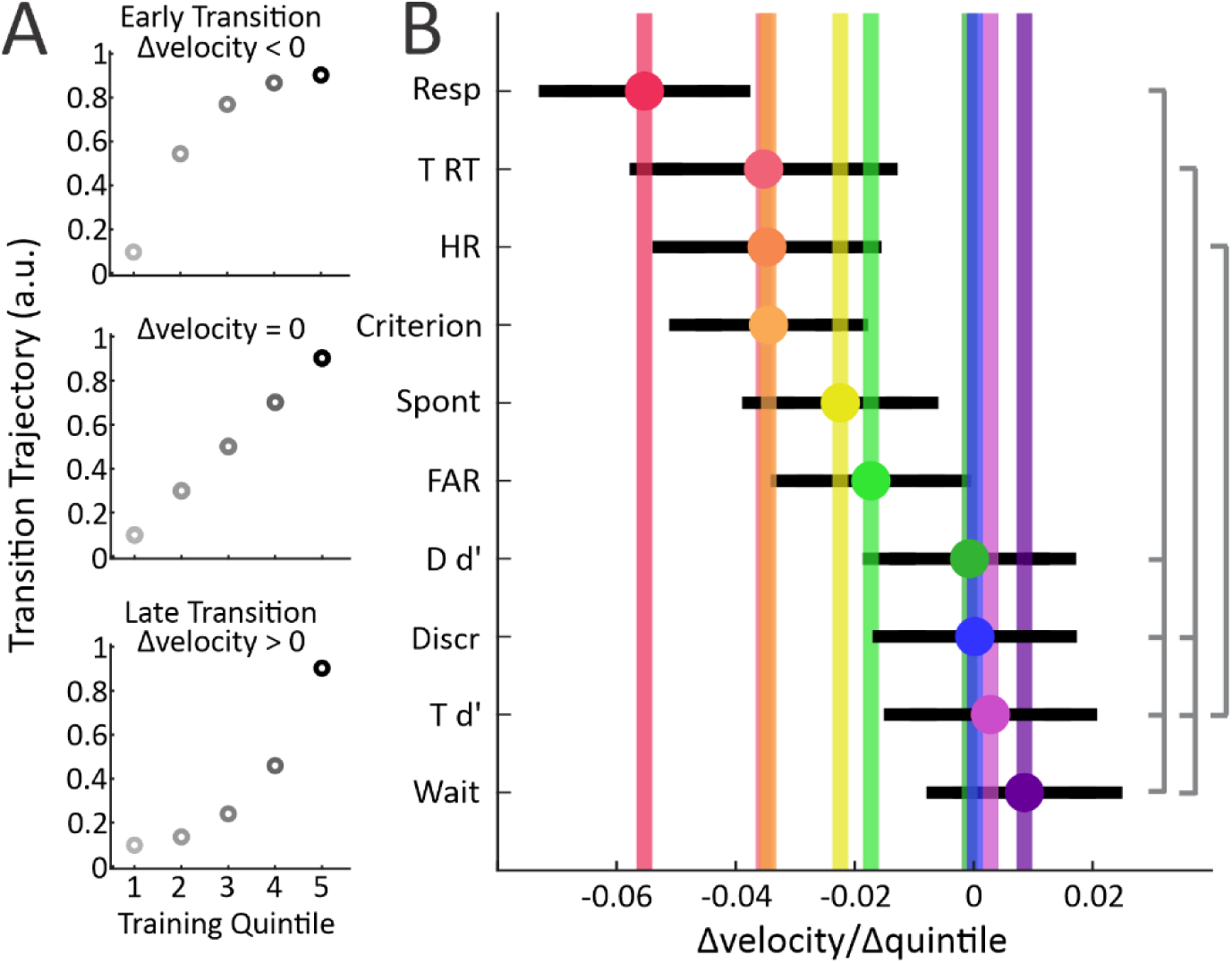
Rate analyses across multiple behavioral measures reveals an order of learning. (A) Illustration of the rate (acceleration versus deceleration) analysis. Top, deceleration (f’’<0) for behavioral measures that change early in training; middle, (f’’=0) for behavioral measures that change throughout training; bottom, acceleration (f’’>0) for behavioral measures that change late in training. (B) Acceleration distributions for phasic behavioral measures across mice, sequenced by increasing acceleration (early changing measures, left; late changing measures, right). Gray lines on the right indicate significant differences (p<0.05) from one-way ANOVA post-hoc multiple comparisons. Resp, response rate; T RT, target reaction time; HR, hit rate; Spont; spontaneous response rate; FAR, false alarm rate; D d’, distractor detection d’; Discr, target vs distractor discrimination d’; T d’, target detection d’; Wait, wait AUC.

### Comparing behavioral measures between mice

While the analyses of Figure 5 capture the average learning dynamics across mice, next we performed behavioral analyses per mouse and compared across the population (Figure 6). We asked whether there was a consistent order of transitioning across these behavioral measures. To assess this, we performed rate analyses of the monophasic and biphasic measures. For each measure and each mouse, we determined the second derivative of the behavioral data (quintile binned and rectified, see methods). We reasoned that early transitioning measures would decelerate [f’’(x) < 0], whereas late transitioning measures would accelerate [f’’(x) > 0] (Figure 6A). These analyses revealed a sequence of transitioning, starting with response rates and ending with wait AUC (one-way repeated measures ANOVA of f’’(x) behavior profiles across mice: F(4.772, 238.6)=4.901, p=0.0004) (Figure 6B). Despite substantial variance across mice, an interesting order of learning emerged. The first four measures reflect post-stimulus responding (response rate, target RT, hit rate, criterion). The next two are the biphasic measures of incorrect (punished) pre-stimulus and post-stimulus responding (spontaneous response rate, false alarm rate). The last four measures include elements of response withholding (distractor detection, target vs distractor discrimination, target detection, wait AUC). We interpret these data as further evidence for multi-staged learning, with the earlier transitioning measures reflecting learning to respond after a whisker stimulus (Fitts/Posner stage 1), and the latter transitioning measures reflecting the shaping of this sensorimotor action to the specific contingencies of this task (Fitts/Posner stage 2).

We also performed rate analyses across mice by sex (n=31 males and 16 females, two-way repeated measures ANOVA within subject: F(45, 405)=7.195, p<0.0001). We found notable sex differences in specific behavioral measures, with male mice displaying earlier transitions in spontaneous response rate, false alarm rate, and wait AUC (from post-hoc pair-wise comparisons, data not shown).

**Figure 7.**
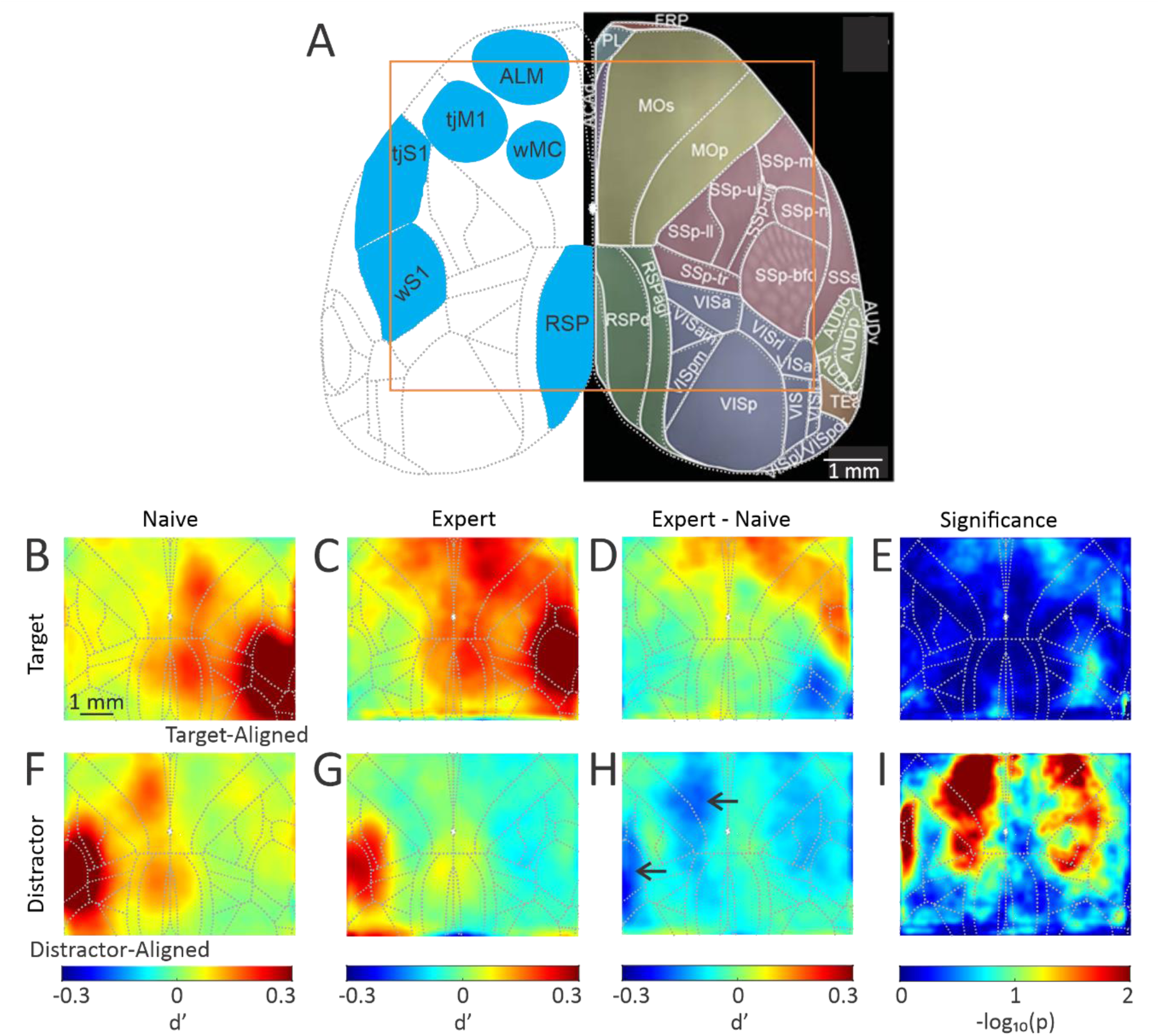
Task learning is associated with unique changes in both target and distractor stimulus-evoked cortical activity. (A) Reference brain (Allen Mouse Brain Common Coordinate Framework) and areas of interest in dorsal neocortex, with the widefield imaging window outlined in orange. (B) Post-stimulus and pre-response cortical activation on target trials in naïve mice. (C) Post-stimulus and pre-response cortical activation on target trials in expert mice. (D) Differences in cortical activation between expert and naïve sessions for target trials [expert – naïve], showing enhanced activations of target-aligned tjSM and bilateral ALM in expert sessions. (F) Post-stimulus and pre-response cortical activation on distractor trials in naïve mice. (G) Post-stimulus and pre-response cortical activation on distractor trials in expert mice. (H) Differences in cortical activation between expert and naïve sessions for distractor trials, showing reduced activations of whisker regions in the distractor-aligned hemisphere (arrows) and parietal and frontal cortices bilaterally. (E and I) Significance maps for D and H, respectively, with the p-value indicated by the colorbar.

### Neuronal correlates of task learning

To determine the cortical modulations associated with task learning, we trained 6 additional mice in the whisker discrimination task and performed longitudinal widefield Ca^2+^ imaging of dorsal neocortex throughout training (Figure 7A). The primary imaging epoch used for the following analyses is 200-300 ms post-stimulus, which includes both post-stimulus and pre-response related signaling. By excluding trials with reaction times <300 ms, these data do not include overt licking or post-reward signaling. In naïve sessions (n=47 sessions), both target and distractor stimuli evoked robust activations of the whisker region of primary somatosensory cortex (wS1), the whisker region of motor cortex (wMC), and the retrosplenial cortex (RSP) (Figure 7B, F). As expected, these activations were mostly unilateral, contralateral to stimulated whiskers. In expert sessions of the same mice (n=20 sessions), the activation patterns changed dramatically and asymmetrically. Target stimuli in expertly trained mice additionally evoked activations of large swaths of frontal cortex (bilateral) and rostral-lateral parietal cortex (unilateral) (Figure 7C). In contrast, distractor stimuli in expertly trained mice evoked mild suppression throughout frontal and parietal cortices, with notably reduced propagation to ipsilateral wMC, ipsilateral RSP, and contralateral wS1 (Figure 7G) (Aruljothi et al., 2020; Zhang & Zagha, 2023).

To quantify these differences, we determined the [expert – naïve] difference maps and performed statistical analyses per pixel across sessions. From these analyses, we find for target stimuli a trend towards increased activation of frontal and parietal regions identified as a pre-motor licking region (anterior lateral motor cortex, ALM) (Guo et al., 2014; Li et al., 2016) and the sensorimotor tongue-jaw region (tjS1 and tjM1, or tjSM) (Mayrhofer et al., 2019) (Figure 7D, E). Interestingly, we did not observe increased activations of whisker-related sensory (wS1) or motor (wMC) regions, as we had anticipated in this whisker discrimination task. In contrast, for distractor trails the [expert – naïve] difference map showed significantly reduced activations of frontal and parietal cortices bilaterally, most robustly in whisker-related regions (Figure 7H, I). Overall, these findings suggest diverse and widespread learning-related changes across dorsal cortex as mice transition from naïve to expert performers.

**Figure 8.**
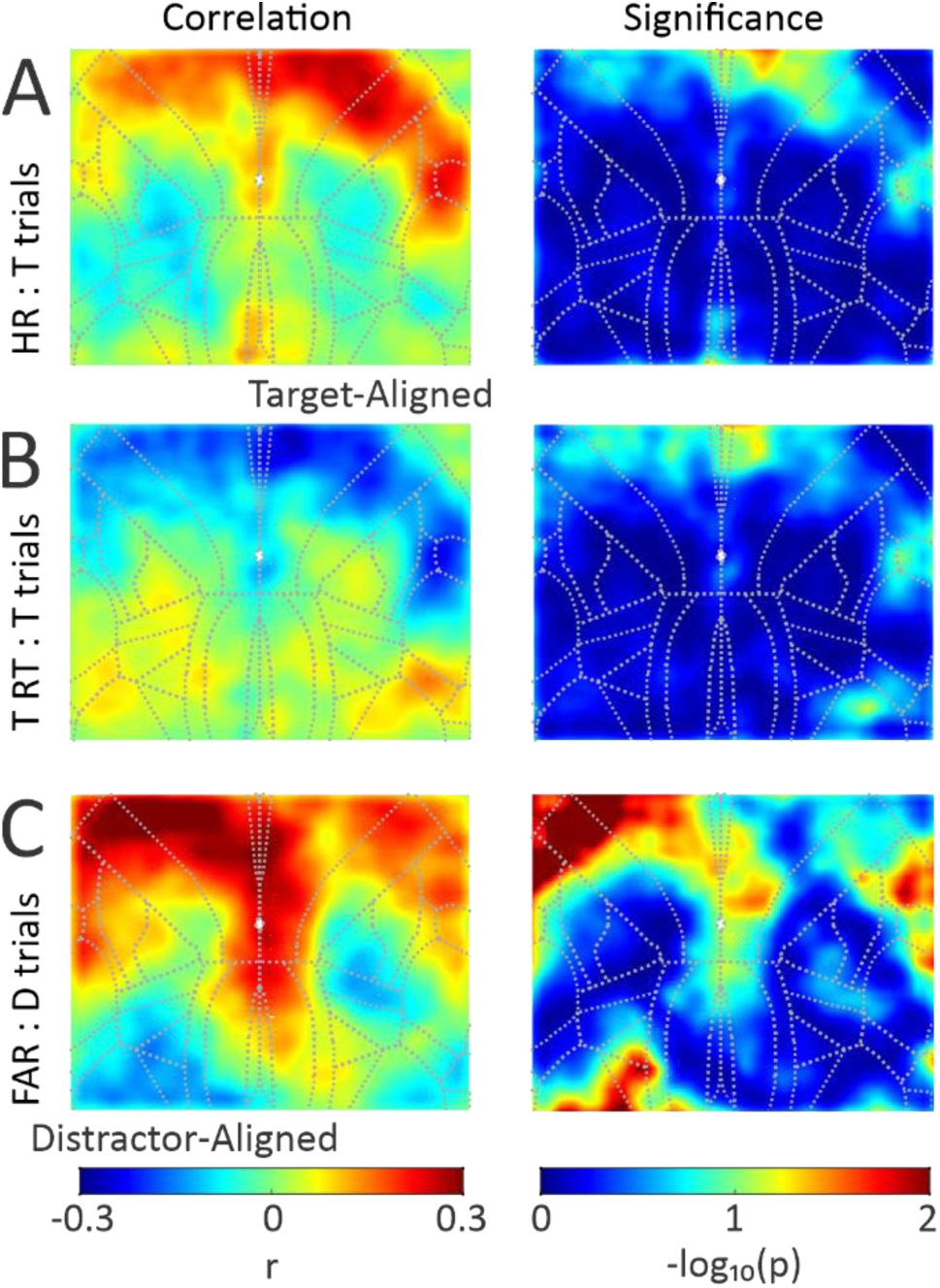
Post-stimulus responding correlates with stimulus-evoked activations of licking-related regions. (A) Left, correlation map of hit rate and target stimulus-evoked cortical activations across training. Right, corresponding significance map. (B) Correlation map (left) and significance map (right) of target reaction times and target stimulus-evoked cortical activations. (C) Correlation map (left) and significance map (right) of false alarm rate and distractor stimulus-evoked cortical activations. Note the consistent correlations between these behavioral measures and activations of licking-related frontal and parietal regions, most prominently in the stimulus-aligned hemisphere.

### Correlations between behavioral measures and neuronal activations

Next, we determined the neuronal correlates of learning across different behavioral measures. We recognize two potential outcomes: 1) all learning measures correlate with the same activation pattern (low dimensional), or 2) different learning measures correlate with different activation patterns (high dimensional). We generated pixel-by-pixel correlation maps between behavioral and neuronal measures across sessions on target or distractor trials, as appropriate. For these analyses, we used all training sessions, from naïve through expert performance. Figure 8A shows the correlation and significance maps between hit rate and neuronal activation on target trials (n=6 mice). We find robust positive correlations between hit rate and activation of unilateral tjSM and bilateral ALM, indicating that increases in hit rate across sessions correlate with increased target-evoked activations of tjSM and ALM. Figure 8B shows the correlation and significance maps for target RT and neuronal activation on target trials. These analyses revealed substantial negative correlations in tjSM and ALM, indicating that reductions in RT across sessions correlate with increased target-evoked activations of tjSM and ALM. The observation that the hit rate and target RT correlation maps are largely overlapping suggests a common neuronal process underlying both learning measures (low dimensional). Notably, we do not observe a consistent correlation between hit rate or target RT and activation of whisker-related sensory or motor regions.

We also determined the correlation between false alarm rate and neuronal activations on distractor trials (Figure 8C). We find similar correlation patterns as with the previous measures, yet more dominant in the distractor-aligned hemisphere. As above, we do not observe a consistent correlation between false alarm rate and activation of whisker-related sensory or motor regions. Altogether, these data suggest a common neuronal correlate driving session-by-session changes in hit rate, target RT, and false alarm rate: stimulus-evoked activations of tongue-jaw and pre-motor licking regions.

**Figure 9.**
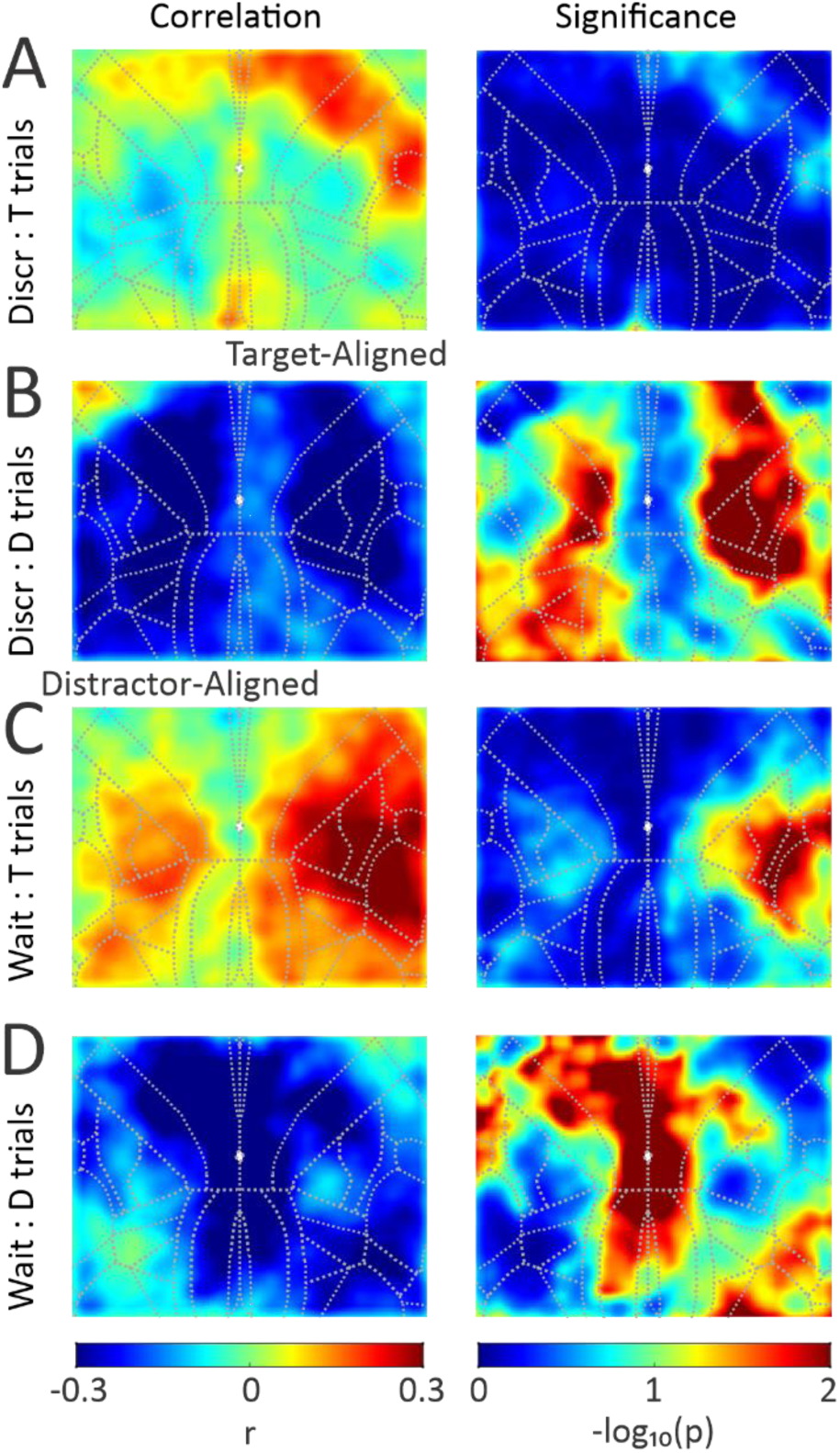
Response withholding correlates with diverse stimulus-evoked activation patterns. (A) Correlation map (left) and significance map (right) of discrimination d’ and target stimulus-evoked cortical activations. (B) Correlation map (left) and significance map (right) of discrimination d’ and distractor stimulus-evoked cortical activations. Note the global negative correlation. (C) Correlation map (left) and significance map (right) of wait AUC and target stimulus-evoked cortical activations. Note the positive correlations within parietal cortices. (D) Correlation map (left) and significance map (right) of wait AUC and distractor stimulus-evoked cortical activations. Note the negative correlations within midline cortices.

### Diversity of behavior-neuronal correlation patterns

The next behavioral measure we investigated was target vs distractor discrimination, the primary outcome measure of this task. When correlating discrimination to neuronal activations on target trials we find the same correlation patterns as for hit rate and reaction time, indicating that increases in discrimination positively correlate with increased activation of tjSM and ALM on target trials (Figure 9A). In contrast, the correlation map between discrimination and neuronal activations on distractor trials shows robust and widespread negative correlation (Figure 9B). This indicates that increases in discrimination negatively correlate with distractor evoked cortical activity. Notably, this pattern on distractor trials is not simply the inverse of the pattern observed on target trials, and therefore suggests a different, more global, neuronal mechanism.

Lastly, we performed correlations between the ability to wait (wait AUC) and target or distractor stimulus activation patterns (Figure 9C-D). These findings were remarkably different than the above patterns. For target trials, increased ability to wait positively correlated with activation of parietal cortex bilaterally, focused on the somatosensory limb regions. In contrast, for distractor trials, increased ability to wait positively correlated with suppression of midline and frontal regions bilaterally. The diverse behavior-neuronal correlation patterns suggest that the different behavioral measures track different underlying neuronal processes (high dimensional). Combined with the behavioral analyses above, our findings support multidimensional learning stages implemented by diverse and widespread neuronal processes during learning of a simple goal-directed task.

## Discussion

In this study, we demonstrate learning-related changes across a host of object-based and temporal-based behavioral measures. In addition to the primary outcome measure (target vs distractor discrimination), these measures capture both overall responding (response rates, criterion) and specific responding to target stimuli (hit rate, hit RT, target detection).

Furthermore, these measures capture two types of inhibitory control: spontaneous withholding during the ITI (spontaneous response rate, wait AUC) and specific withholding to distractor stimuli (false alarm rate, false alarm RT, distractor detection). Across 52 mice, changes in these measures suggest an ordered learning process, with increased responding occurring early in training and strategic withholding occurring later in training. Furthermore, our behavior-neuronal comparisons suggest different neuronal mechanisms underlying these behavioral changes. Responding to target and distractor stimuli correlated with stimulus-evoked activations of response (lick)-related sensorimotor cortices. Neuronal correlates of withholding were more diverse, and unique for each specific behavioral measure. Altogether, our studies begin to reveal the multi-staged learning of this goal-directed task, and the diverse and widespread neuronal changes associated with each learning stage.

We propose adopting the Fitts/Posner framework (Fitts & Posner, 1967) as a general structure of mouse goal-directed task learning. We identify three important reasons for adopting this framework. First, if there is an orderly progression of learning, then identifying the learning stages will aid in tracking the learning process. Second, and relatedly, mouse models of neuropsychiatric disease show learning deficits in goal-directed tasks (Goel et al., 2018; Hölter et al., 2015; Huynh et al., 2009; Remmelink et al., 2016). Localizing the learning deficits to specific learning stages is essential for understanding the underlying neuronal mechanisms involved. For example, a mouse model with impairments at Fitts/Posner stage 1 (learning the required sensorimotor action) is likely to have a very different pathophysiology than a mouse model with impairments at Fitts/Posner stage 2 (refining the learned action according to the task rules). Further distinctions can be made within a learning stage from specific behavioral measures. For example, poor performance due to impaired spontaneous withholding (measured in our task by wait AUC) is likely to have a very different pathophysiology than poor performance due to impaired distractor stimulus withholding (measured in our task by false alarm rate and distractor detection). Third, a growing number of mouse studies are aimed at revealing the neuronal mechanisms underlying task learning (e.g., (Audette et al., 2019; Huber et al., 2012; Lacefield et al., 2019; Le Merre et al., 2018; Makino et al., 2017; Ray et al., 2023; Roy et al., 2021)). Correlating the changes in specific behavioral and neuronal measures will vastly improve our ability to identify the neuronal mechanisms underlying specific learning processes, and make more precise predictions about the effects of causal neuronal manipulations on learning and performance outcomes. This includes progression to Fitts/Posner stage 3, the autonomous stage requiring less cognitive effort, which may explain the disengagement of neocortex after task acquisition (Hwang et al., 2021; Kawai et al., 2015).

Our behavioral monitoring approach is complimentary to behavioral modeling studies that track performance across learning (Roy et al., 2021). Additionally, discrete state models (Ashwood et al., 2022; Pisupati et al., 2021) are being used to identify fluctuations in behavioral strategy. These types of analyses can be applied to multi-staged learning. However, models of behavioral learning and performance are limited to the strategies defined in the model. In this study, we do not predefine strategies across learning, but instead analyze a host of behavioral measures and use their dynamics across learning to infer strategies. Our model-free approach may reveal additional strategies not considered in existing behavioral models, and therefore can be used to guide model design.

Another insight from this study is the importance of considering temporal processes in object-based tasks. Our ITI distributions were intended to reduce temporal expectancy of target stimulus delivery (Coull et al., 2011; Fiorillo et al., 2008; Ghose & Maunsell, 2002; Nobre et al., 2007; Zariwala et al., 2013). Nevertheless, there remains temporal regularity in the structure of all behavioral tasks. Subjects may learn these temporal regularities to improve task performance by predicting windows of opportunity in task structure (Reyes et al., 2020; Toda et al., 2017). Furthermore, subjects may exploit the temporal regularities to solve a task in a way that was unspecified (Kawai et al., 2015). By considering temporal processes in object-based tasks, researchers may reveal understudied task dimensions that are critical for optimal performance and likely contribute to behavioral dysfunctions in neuropsychiatric disease (Emmons et al., 2017; Grondin, 2010; Toplak et al., 2006).

Longitudinal widefield imaging of mouse neocortex from naïve through expert task performance revealed diverse neuronal correlates of task learning (Figure 7). Using correlation analyses, we attempted to further relate these neuronal changes to specific behavioral processes. Interestingly, changes in post-stimulus responding (for target and distractor stimuli) correlated with stimulus-evoked activation of licking-related ALM and tongue-jaw sensorimotor cortices (Figure 8). This is consistent with activations of these regions in other goal-directed tasks with licking read-outs (e.g., (Chen et al., 2017; Mayrhofer et al., 2019; Musall et al., 2019; Salkoff et al., 2020)). Surprisingly, post-stimulus responding was not correlated with stimulus-evoked activations of the relevant sensory regions (wS1). It is possible that the specific stimuli used in this task, well separated across whisker fields, does not require robust functional plasticity of wS1 processing. If so, then robust plasticity would be expected only downstream of wS1, in linking the whisker stimulus to the licking response (Petersen, 2019). And yet, other studies have shown learning-related changes in wS1 sensory processing (Chen et al., 2015), including for a whisker detection task (Sachidhanandam et al., 2013). Given the course resolution of our widefield imaging method (sampling from the superficial processes of populations of neurons), we cannot rule out important changes in wS1 signaling at the level of individual neurons and specific neuronal sub-types. Nonetheless, our findings suggest that the most robust neuronal changes underlying post-stimulus responding are downstream of wS1.

Lastly, our behavior-neuronal analyses provide further evidence for the multidimensional nature of inhibitory control (Robbins & Dalley, 2017; Robbins et al., 2012). We infer the neuronal correlates of withholding to distractor stimuli by the correlation between discrimination and distractor evoked activations (Figure 9B). Interestingly, we find that this is not simply the inverse of post-stimulus responding (localized to licking-related frontal and parietal regions), but is distributed globally throughout dorsal neocortex. These data suggest a global reduction in distractor stimulus-evoked activation, possibly signaled by broadly-projecting neuromodulatory systems. Furthermore, the neuronal correlates of spontaneous withholding (Figures 9C and 9D) differed dramatically from distractor stimulus withholding (Figure 9B). Additional recording studies from cortical and sub-cortical regions, correlated with causal manipulations, are required to further reveal the neuronal mechanisms underlying the behavioral and cognitive processes identified in this study.

## Acknowledgements

This work was supported by the National Institutes of Health (R01NS107599 to E.Z.). We thank Drs. Hongdian Yang and Anubhuti Goel for many helpful discussions throughout the project. We thank Angelina Lam and Drs. Zhaoran Zhang, Behzad Zareian, and Manas Kinra for valuable critique and collegial support.

